# Ectopic Reconstitution of a Spine-Apparatus Like Structure Provides Insight into Mechanisms Underlying Its Formation

**DOI:** 10.1101/2024.04.16.589782

**Authors:** Hanieh Falahati, Yumei Wu, Pietro De Camilli

**Affiliations:** HHMI; Departments of Neuroscience and Cell Biology; Program in Cellular Neuroscience, Neurodegeneration, and Repair, Yale University School of Medicine, 100 College Street, New Haven, 06511, CT, USA

**Keywords:** organelle specialization, neuronal ER, cytoskeleton

## Abstract

The endoplasmic reticulum (ER) is a continuous cellular endomembrane network that displays focal specializations. Most notable examples of such specializations include the spine apparatus of neuronal dendrites, and the cisternal organelle of axonal initial segments. Both organelles exhibit stacks of smooth ER sheets with a narrow lumen and interconnected by a dense protein matrix. The actin-binding protein synaptopodin is required for their formation. Here, we report that expression in non-neuronal cells of a synaptopodin construct targeted to the ER is sufficient to generate stacked ER cisterns resembling the spine apparatus with molecular properties distinct from the surrounding ER. Cisterns within these stacks are connected to each other by an actin-based matrix that contains proteins also found at the spine apparatus of neuronal spines. These findings reveal a critical role of a synaptopodin-dependent actin matrix in generating cisternal stacks. These ectopically generated structures provide insight into spine apparatus morphogenesis.

## Introduction

The endoplasmic reticulum (ER) is represented by an interconnected endomembrane system which extends into all regions of the cell cytoplasm [1–3]. However, it is organized in morphological and functional distinct domains, more so in specialized cells such as neurons [3–10]. A most notable example of specialized ER subdomain is the spine apparatus, strategically positioned in dendritic spines near synapses [11, 12]. This structure is represented by a stack of interconnected flat smooth ER cisterns with a narrow lumen, which are separated from each other by a dense proteinaceous matrix and are connected to the bulk of the ER via a narrow tubule that travel along the spine’s stalk [3, 12–14]. A morphologically similar ER specialization, called the cisternal organelle, can be observed at axonal initial segments [15, 16]. Little is known about the functions of these ER subdomains, although there is evidence that the spine apparatus may have a role in synaptic plasticity [17–22]. A function in Ca^2+^ storage and Ca^2+^ signaling has been proposed [21, 23, 24], although the contribution of its peculiar shape to such function remains unclear. Alterations in the morphology of the spine apparatus have been reported in response to long-term potentiation [25] and pathological conditions [26–29]. While a number of studies have investigated the morphogenesis of ER subdomains like rough ER sheets and smooth tubules [30–33], mechanisms underlying the formation of the spine apparatus and the cisternal organelle remain elusive.

Previous reports have shown that synaptopodin, a protein selectively concentrated in the spine apparatus and cisternal organelle in the brain, is required for the formation of these structures [17]. Synaptopodin is an actin-binding protein and, accordingly, the spine apparatus and cisternal organelle are localized in actin rich regions of dendritic spines and axonal initial segments [14, 34–37]. Moreover, to discover other proteins associated with these structures, we had carried out a search of protein neighbors of synaptopodin using in vivo proximity proteomics and this approach identified several actin-related proteins [14]. These findings suggested a major role of actin and an actin-related machinery in the generation of the spine apparatus.

Formation of the spine apparatus and cisternal organelle involves several steps. Synaptopodin, which lacks a transmembrane region, must associate directly or indirectly with the ER. The ER, which typically consists of tubules in dendrites, must expand into sheets. These ER sheets need to be stacked on top of each other via a dense intervening matrix and this process must correlate with a narrowing of their lumen. The contributions of synaptopodin and synaptopodin neighbors to these steps, remain open questions in neuronal cell biology.

Goal of this study was to begin addressing these questions. We show that the spine apparatus and synaptopodin are conserved from fly to humans, and that a highly conserved region of this protein is necessary for its association with ER. We reveal a dual role of synaptopodin in generating actin bundles and in linking them to the ER. In addition, we show that expression in COS-7 cells of a synaptopodin fusion protein directly anchored to the ER is sufficient to induce the formation of spine apparatus-like structures. Our findings suggest that interactions between synaptopodin and an actin-rich cytomatrix drive the formation of these organelles. Ectopic spine apparatus-like structures generated in non-neuronal cells represent a powerful model system to gain further insight into mechanisms that generate these specialized and poorly understood ER-based organelles.

## Results

### Synaptopodin can crosslink ER to actin filaments in neurons

The two neuron-specific ER specializations positive for synaptopodin, the spine apparatus and the cisternal organelle, share special morphological features. They are composed of stacks of smooth ER sheets which have a very narrow lumen and are connected to each other by a proteinaceous dense matrix (Fig 1A-B)[11, 12]. Concentrations of both endogenous and exogenous synaptopodin - as revealed by immunofluorescence and by expression of fluorescently tagged synaptopodin respectively - are also observed in dendritic spines and axonal initial segments in cultured mouse hippocampal neurons [14, 35] (Fig. 1C, S1A-C). Fluorescent exogenous (and thus overexpressed) synaptopodin also accumulated in spots or elongated structures (up to several micron long) in dendritic shafts and perikarya (Fig. 1C). The nature of these synaptopodin accumulations was analyzed by EM. To this aim, cultured DIV7 neurons were transduced with AAV2/9 viruses encoding EGFP-synaptopodin and cells showing high abundance of these fluorescent structures were selected for analysis by correlative light and electron microscopy (CLEM). EM observation revealed that the synaptopodin-positive accumulations were represented by bundles of filaments with the expected morphology of F-actin sandwiched between ER cisterns (Fig. 1D, S2). In some cases, only a thin layer of actin separated ER cisterns, thus generating structures reminiscent of ER stacks of the spine apparatus. These results show that synaptopodin, when expressed in neurons, not only can bundle F-actin, as shown previously [14, 34, 36], but can also crosslink F-actin bundles to the ER. They support a key role of synaptopodin in the formation of the spine apparatus and cisternal organelles, which are ER-based structures.

**Fig. 1.**
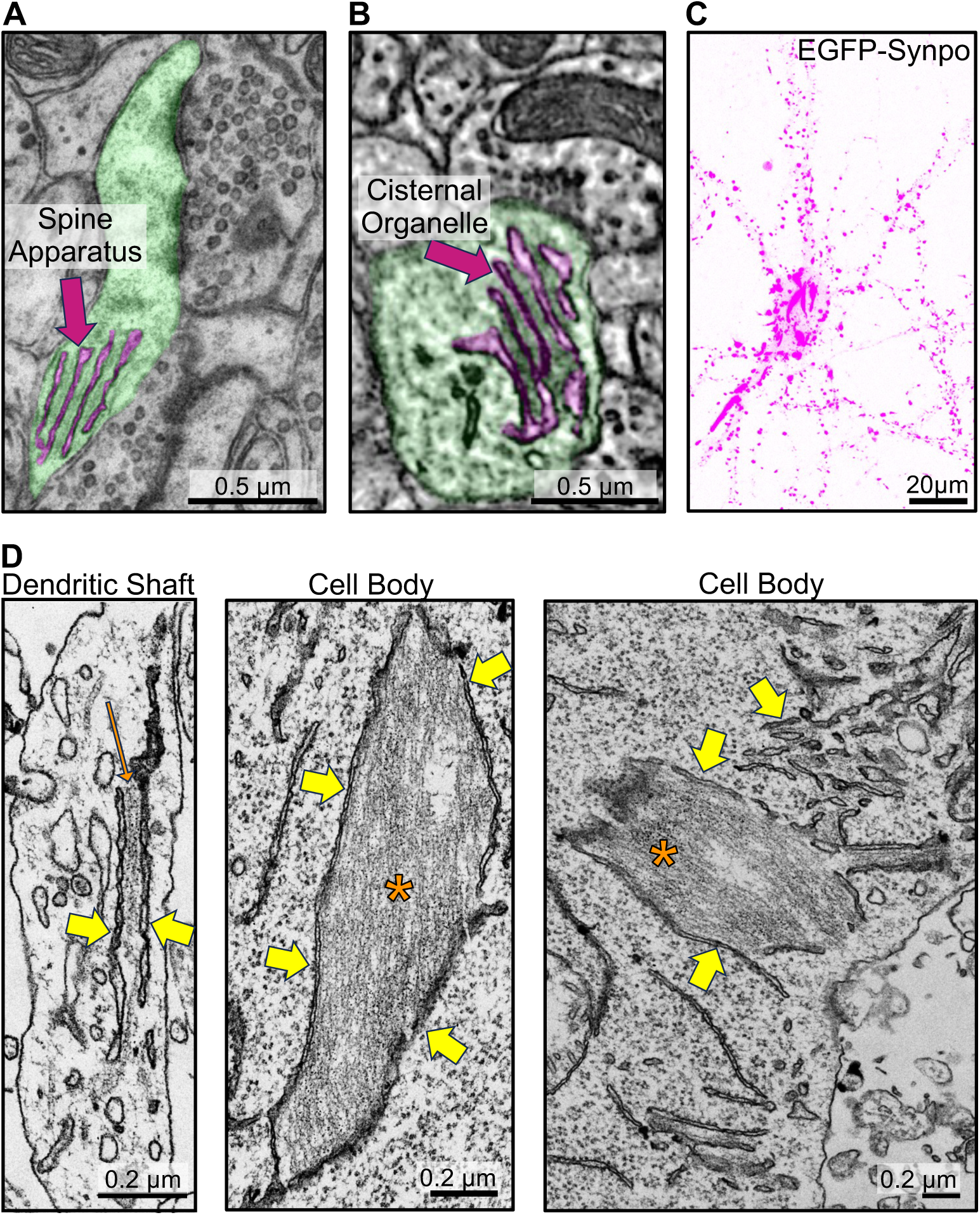
Synaptopodin mediated recruitment of actin filaments to the ER in neurons. **A.** Transmission electron microscopy (TEM) image of a pseudocolored dendritic spine (green) with spine apparatus (magenta). **B**. Serial block-face scanning electron microscopy (SBF-SEM) image of a pseudocolored cisternal organelle (magenta) at an axonal initial segment (green) (image from microns-explorer.org)[38]. **C–D**. Fluorescent image (**C**), and TEM (**D**) of a cultured mouse hippocampal neuron infected with AAV2/9 encoding EGFP-synaptopodin. Note in **C**large accumulations of synaptopodin in the central region of the cells in addition to the punctate fluorescence in dendrites. Many of smaller puncta along dendrites correspond to the expected accumulations of synaptopodin in the spine apparatus of dendritic spines. Yellow arrows in **D** point to the ER and one orange arrow and asterisks point to actin bundles.

### Synaptopodin can crosslinks ER to F-actin bundles also in fibroblasts

We further investigated the relationship between synaptopodin, actin and the ER by expressing fluorescently tagged synaptopodin (mRFP- or EGFP-synaptopodin) in COS-7 cells, a fibroblastic cell line whose flat morphology facilitates the analysis of these relationships by fluorescence microscopy. As reported [14, 34], strong colocalization of mRFP-synaptopodin with F-actin (phalloidin staining) was observed in these cells (Fig. 2A). A subset of these synaptopodin- and actin-positive elements had the appearance of stress fibers. Another subset of them were thick linear inclusions generally located deep in the cytoplasm and away from the plasma membrane (Fig. 2A; see color coding of depth in 2B). At EM observation these structures were found to correspond to bundles of actin sandwiched between tightly apposed ER cisterns (Fig. 2C) similar to those observed in neurons overexpressing synaptopodin (Fig. 1D). No such structures were observed in non-transfected COS-7 cells.

**Fig. 2.**
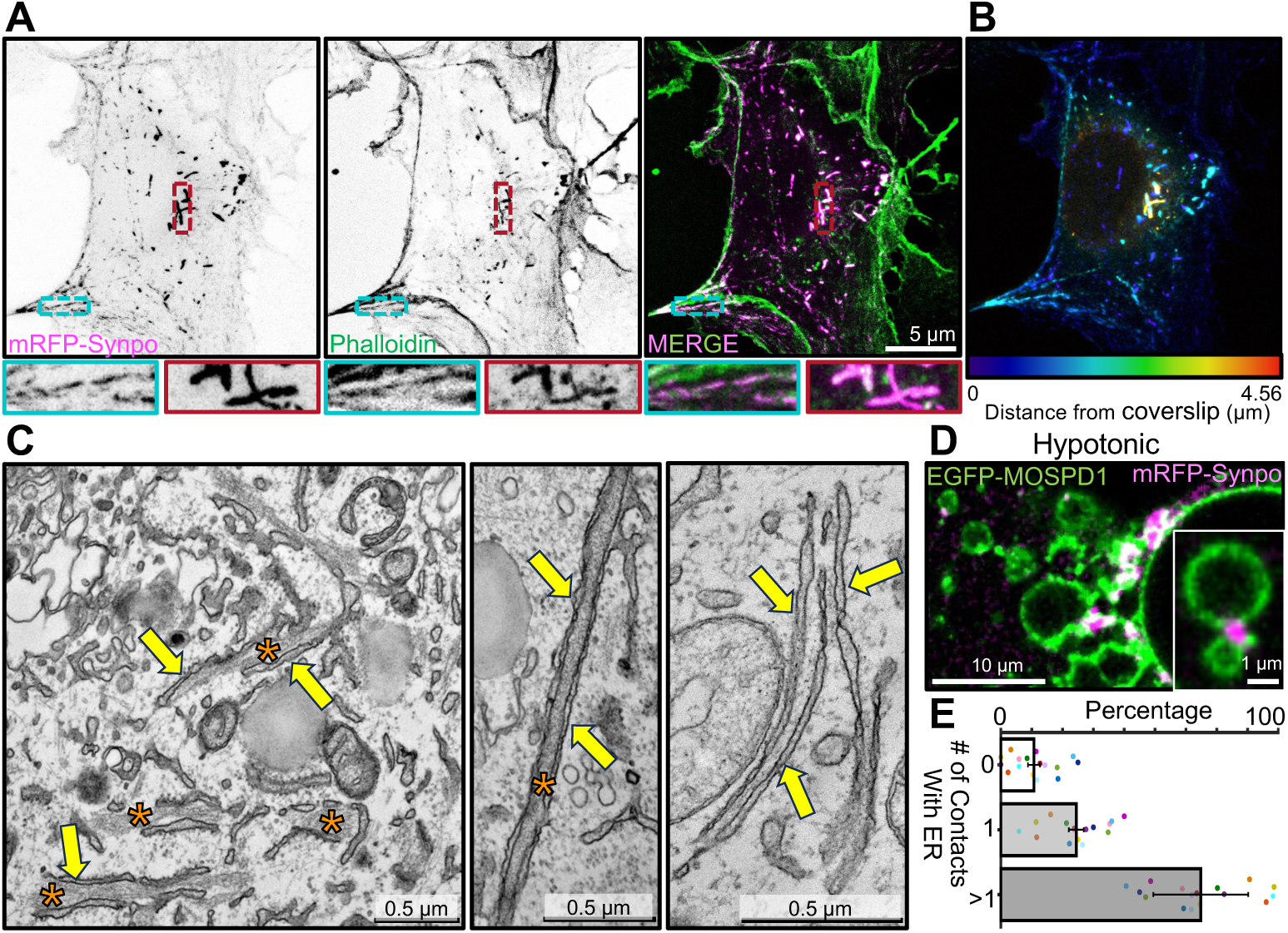
Localization of exogenous synaptopodin in COS-7 cells both on stress fibers and on actin-rich structures associated with ER. **A**. COS-7 cells expressing mRFP-synaptopodin displaying strong overlap of the mRFP fluorescence with phalloidin staining. High magnification views of a stress fiber (blue) and a non-stress fiber linear assembly (red) are shown at the bottom. **B**. mRFP-synaptopodin signal from **A**is shown with color-coding based on the distance from the coverslip. Stress fibers are closer to coverslip while non-stress fiber actin assemblies are present anywhere within the cell. **C**. TEM of a cell expressing mRFP-synaptopodin showing actin bundles (orange asterisks) sandwiched between ER sheets (yellow arrows). **D**. COS-7 cells expressing mRFP-synaptopodin (magenta) and the ER protein EGFP-MOSPD1 (green) were exposed to hypotonic conditions. The accumulation of synaptopodin at the interface between ER elements reveals its direct or indirect association with the ER membrane. **E**. Percentage of mRFP-synaptopodin puncta with or without association with ER membrane in COS-7 cells exposed to hypotonic conditions is shown. Data from different cells are shown as a dot with different color.

The tight attachment of these synaptopodin-positive elements to the ER was further supported by acute exposure of cells to drastic hypotonic conditions, an approach previously used to examine ER contacts with other structures [39, 40]. Upon hypotonic shock, the ER rapidly vesiculates, but many of its contacts with other structures persist for some time and can be visualized as the cell swells and undergoes lysis. For these experiments cells were transfected not only with mRFP-synaptopodin, but also with EGFP-MOSPD1 as a marker of the ER [41] (Fig. 2D, Video 1). After 60 minutes from the beginning of the hypotonic shock, 90% (*±* 2%) of the synaptopodin positive elements were still in contact with the ER, confirming the occurrence of a link between them (Fig. 2E). Moreover, 65% (*±* 15%) of them were at contacts between ER vesicles consistent with their role in crosslinking ER cisterns before the hypotonic lysis.

We conclude that even in fibroblasts synaptopodin not only can bundle actin, but can also link such bundles to the ER, implying the presence in this protein of sites that directly or indirectly bind the ER membrane.

### Synaptopodin has multiple actin binding sites

Synaptopodin is not predicted to have major folded domains (Fig. 3A). In order to identify the region(s) within synaptopodin responsible for its property to crosslink actin bundles to the ER, we generated several truncation mutants of this protein and tested them for their property to generate the actin-rich ER-associated structures described above. Previous work showed that residues 384-473 include the actin binding region of synaptopodin [42] (Fig. 3B). We found that deletion of residues 384-473 did not eliminate its property to associate with F-actin, suggesting the presence of additional actin binding sites (Fig. 3C-D). However, this mutant no longer formed the F-actin positive inclusion that reflect the ER-associated actin bundles (see Fig. 2A). Both regions upstream (residues 1-384) or downstream (residues 474-690) retained their ability to localize to stress fibers (Fig. 3E-F), confirming that synaptopodin can directly or indirectly interact with F-actin through multiple regions.

**Fig. 3.**
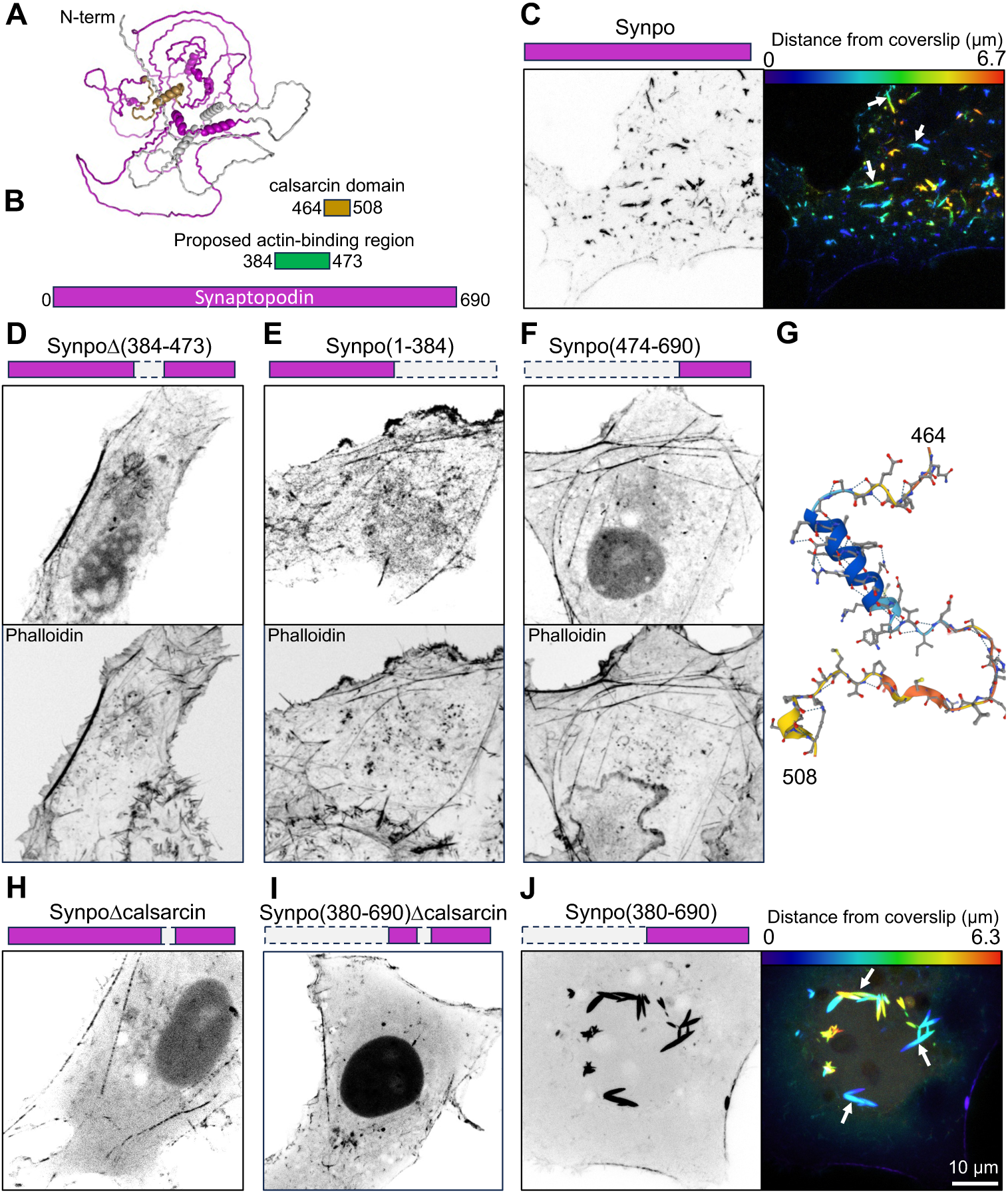
Synaptopodin regions involved with F-actin and ER binding. **A**. Predicted structure of mouse synaptopodin (accession number: Q8CC35-1) based on Alphafold. The synaptopodin construct used for this study is the short splice variant (accession number: Q8CC35-3) that lacks the first 239 residues (shown in gray) and is the brain-specific isoform. The calsarcin domain is shown in gold. **B.** Domain representation of synaptopodin, showing in green the previously reported actin binding region [42] and its partial overlap with the calsarcin domain (gold). **C**. COS-7 cell expressing fluorescently tagged full-length synaptopodin and showing the prominent accumulation of synaptopodin inclusions (arrows) that represent actin-ER assemblies. The synaptopodin signal is shown in gray scale (left), and color-coded based on distance from coverslip (right). **D–F**. COS-7 cells expressing the indicated deletion constructs of synaptopodin (magenta color) lacking the reported actin binding region showing that all of them still partially colocalize with F-actin as indicated by phalloidin staining. **G**. Predicted structure of the calsarcin region of synaptopodin based on Alphafold. **H** and **I**. COS-7 cells expressing synaptopodin deletion constructs showing that the construct including the calsarcin domain (**J**), but not the ones excluding this domain (**H** and **I**) induce the formation of synaptopodin inclusions (arrows) that represent actin-ER assemblies (see Fig. 2).

We observed that the fragment of synaptopodin previously identified as the actin binding region [42] and found by us to be required for the formation of ER-associated inclusions, i.e. the fragment comprising a.a. 384-473, partially overlapped with a 45 a.a. stretch classified as calsarcin domain in Pfam (a.a. 464-508) (Fig. 3B, G). Deletion of these residues from full length synaptopodin or its C-terminal fragment (a.a. 380-690) abolished the F-actin positive inclusions associated with ER (Fig. 3H-I), while such inclusions were prominent in cells expressing the C-terminal fragment comprising this region (Fig. 3J). However, fusing fragment 380-508 of synaptopodin (which comprises these 45 a.a.) to the C-terminal of mRFP was cytosolic, indicating that the calsarcin domain of synaptopodin is not sufficient to anchor a protein to the ER. Thus, the precise nature of the interaction of synaptopodin with the ER remains to be elucidated.

The region of synaptopodin defined in databases as calsarcin domain comprises an alpha-helix followed by a hairpin (Fig. 3G). The alpha-helix share similarity to an alpha-helix of myozenin [43] (also known as calsarcin, hence the name of the domain) (Fig. S3A). Interestingly, myozenin, like synaptopodin, binds Pdlim family proteins [44], which include Pdlim7, a synaptopodin interactor [14], although myozenin does not use the calsarcin domain for such binding [44]. The calsarcin domain is also highly conserved in the synaptopodin paralogues synaptopodin 2 and synaptopodin 2-like [45], and extends to the hairpin region (67% similarity, Fig. S3A). Moreover, when expressed in COS-7 cells, fluorescently tagged synaptopodin 2-like protein also localizes on internal F-actin inclusions similar to those generated by synaptopodin (Fig. S3B). Based on public gene expression databases (e.g. https://tabula-sapiens-portal.ds.czbiohub.org), synaptopodin 2 and synaptopodin 2-like have a similar expression pattern as synaptopodin in human brain, but are mainly absent from mouse neurons. This likely explains why in mice the lack of synaptopodin is sufficient to abolish presence of spine apparatus and cisternal organelles.

### Synaptopodin and the spine apparatus are conserved from fly to humans

To delve deeper into the contribution of synaptopodin to spine apparatus formation, we explored its occurrence throughout evolution. Analysis of the evolutionary tree of synaptopodin using Panther [47] revealed that one of its early orthologues is CG1674 in *Drosophila melanogaster*. CG1674 is an actin binding protein [48], and its calsarcin domain shares 47% similarity with that of human synaptopodin (Fig. 4A). Accordingly, expression of mRFP-CG1674 in COS-7 cells resulted in the formation of F-actin inclusions localized deep in the cytoplasm similar to those observed upon expression of mammalian synaptopodin (Fig. 4B–C). Moreover, coexpression of mRFP-CG1674 with mouse synaptopodin (EGFP-synaptopodin) resulted in their precise colocalization (Fig. S3C). Since the presence of spine apparatus was only reported in mammals, we searched the entire brain electron microscopy dataset of *D. melanogaster*, FlyWire [46]. Interestingly, while most *D. melanogaster* neurons lack spines, a subset of visual system interneurons called LPTCs have actin-rich dendritic spines similar to that of vertebrates [49] (Fig. 4C) and in such spines the presence of two to three ER sheet elements closely apposed to each other by an intervening proteinaceous density can be observed (Fig. 4D–E). While such ER elements have a wide lumen, in contrast to ER elements of the mammalian spine apparatus, it remains possible, that such width may reflect a dilation due to fixation conditions.

**Fig. 4.**
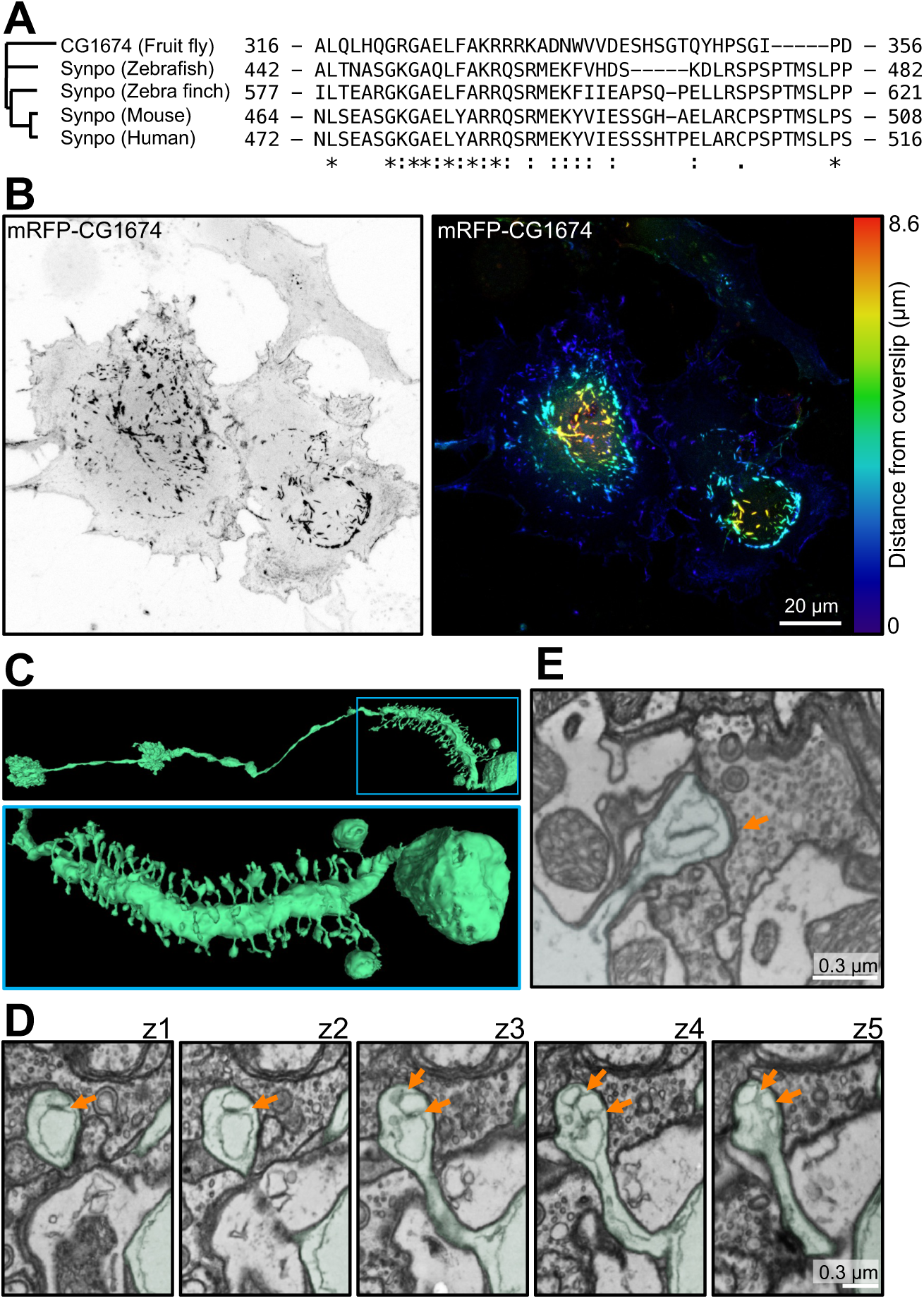
Presence of synaptopodin and spine apparatus in *Drosophila melanogaster*. **A**. Residues 464-508 of mouse synaptopodin (calsarcin domain) are highly conserved among synaptopodin orthologues in vertebrates and *D. melanogaster* orthologue, CG1674. **B**. COS-7 cell expressing the fluorescently tagged *D. melanogaster* synaptopodin orthologue. A prominent accumulation of CG1674 inclusions, similar to those observed in cells expressing fluorescently tagged mouse synaptopodin (See Fig. 2 and 3C), is visible. The CG1674 signal is shown in gray scale (left), and color-coded based on distance from coverslip (right). **C**. *D. melanogaster*’s LPTC neuron reconstructed from 3D SBF-SEM images (from FlyWire [46]) showing spines along its major process. Low and high magnification views are shown at the top and bottom, respectively. **D**. SBF-SEM sequential optical sections from a spine of the neurons shown in **C**, revealing ER elements closely apposed via an intervening density (orange arrows) in the spine head. This structure is reminiscent of a spine apparatus with dilated ER cisterns, possibly due to preparation artifacts. **E**. Another example of a dendritic spine of *D. melanogaster* containing a structure reminiscent of a spine apparatus, but with a dilated ER.

Together, these observations raise the possibility that synaptopodin and the spine apparatus may be conserved from flies to humans.

### A Spine-Apparatus-Like structure (SAL) is reconstituted in COS-7 cells

The results and observations reported above reveal that synaptopodin can bind and bundle actin and also link actin to the ER. These properties suggest that synaptopodin could mediate the close apposition of ER elements observed in the spine apparatus via an intervening actin matrix. However, a major difference observed between the spine apparatus and the structures induced by synaptopodin overexpression in both neurons and COS-7 cells is the thickness of the actin bundles separating the ER cisterns. Such greater thickness is most likely due to actin crosslinking by synaptopodin not in contact with the ER. What limits actin bundling in dendritic spines remains unclear. We explored what would happen if we restricted synaptopodin binding and bundling of F-actin exclusively to when it is in contact with the ER, by anchoring it to the ER via a transmembrane region. This was achieved by using a construct (synaptopodin-ER, Fig. 5A) in which synaptopodin (as a fluorescent fusion protein) is fused to the N-terminus of Sec61*β*, an ER resident protein anchored to this organelle through a C-terminal transmembrane region [50]. As shown previously [14], expression of this chimera in COS-7 cells resulted in the formation of elongated inclusions (Fig. 5B–E) that were positive for actin - as shown by phalloidin staining (Fig. 5D)- and bore a resemblance to those induced in the cell bodies of neurons (Fig. 1C) and COS-7 cells (Fig. 2A) by synaptopodin overexpression. In cells with high levels of synaptopodin-ER such structures were extremely abundant (Fig. 5B and E).

**Fig. 5.**
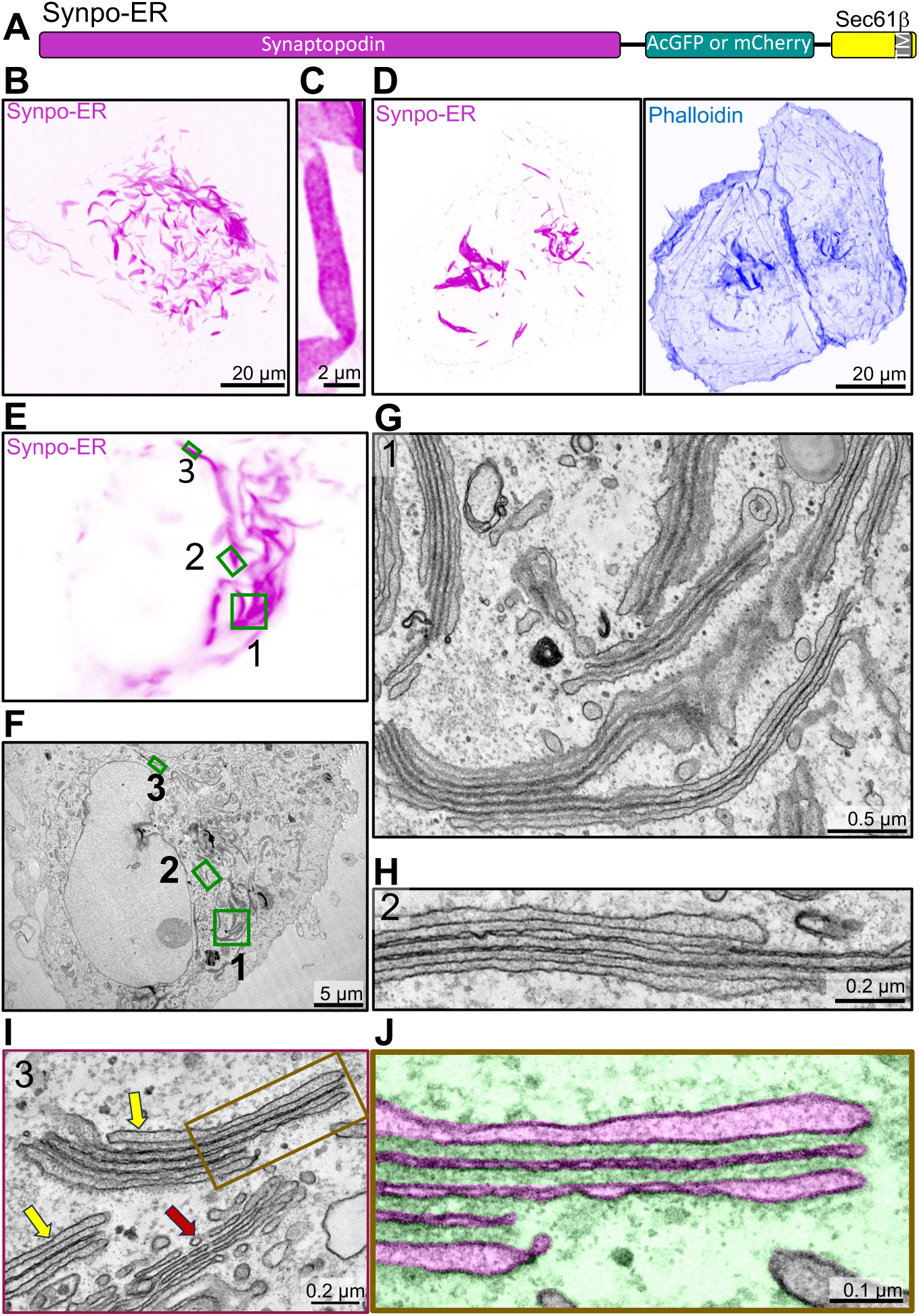
Generation of spine apparatus-like structures (SALs) in COS-7 cells. **A**. Diagram of the synaptopodin construct anchored to the ER (Synpo-ER) by its fusion to Sec61*β*, which is embedded in the ER by a C-terminal transmembrane region. **B**. Confocal image of a COS-7 cell expressing Synpo-ER. **C**. High magnification view of a SAL as imaged by an AiryScan confocal microscope. **D**. Presence of F-actin in SALs as indicated by phalloidin staining. **E** and **F**. Light microscope image (**E**) of a COS-7 cell expressing synaptopodin with its corresponding EM image (**F**). High magnification EM images of the regions 1 - 3 framed by green rectangles in fields **E**and **F** are shown in **G–I**, respectively. Moreover, a portion of the SAL in **I** is shown at higher magnification in **J**with the ER lumen pseudocolored in magenta and the cytosolic space in green. SALs are represented by ER stacks with morphological features similar to those of the spine apparatus and of the cisternal organelle. Red and yellow arrows in **I** point to the Golgi complex and to two SALs, respectively.

Strikingly, CLEM of synaptopodin-ER expressing cells (Fig. 5E–J) showed that the fluorescent signal of synaptopodin-ER corresponded to ER sheets stacked together with only a thin dense matrix separating them (Fig. 5G–J, S4A–B). These stacks, as observed by FIB-SEM, often extended over several micrometers in length (Fig. S4C and Video 2). FIB-SEM also showed that cisterns were continuous with the tubular ER (Video 2). Within these stacks, ER sheets had a very narrow, nearly absent lumen, with the exception of the first and last cisterns. Accordingly, DsRed-KDEL was nearly excluded from these stacks in live cells (Fig. S5), implying that most luminal ER proteins are likely excluded from these structures. Overall, the appearance of these ER stacks, which we will refer to henceforth a SALs (Spine Apparatus-Like) had key distinctive morphological features of the spine apparatus and cisternal organelle. Moreover, as in the spine apparatus, a dense matrix was observed between ER sheets (Fig. 1A–B), although the spacing between the sheets (28.2 *±* 0.8 nm) (Fig. S4D) was narrower than in the spine apparatus (51 *±* 2 nm) (Fig. S4D). Interestingly, housekeeping ER membrane proteins such as VAP-B and STIM1 are also excluded from these structures (Fig. S5) suggesting that while SALs are continuous with the rest of ER, they are molecularly distinct subdomains. In addition to actin (Fig. 6A), the proteins that we previously identified to localize to the spine apparatus in the brain, *α*-actinin 2, Pdlim7, Magi1, and Magi2, all localize to SALs [14](see also Fig. 6B-C, S5), confirming that SALs share molecular components with the spine apparatus. We conclude that expression of synaptopodin-ER generates ER specializations that can provide insight into mechanisms underlying formation of the spine apparatus.

**Fig. 6.**
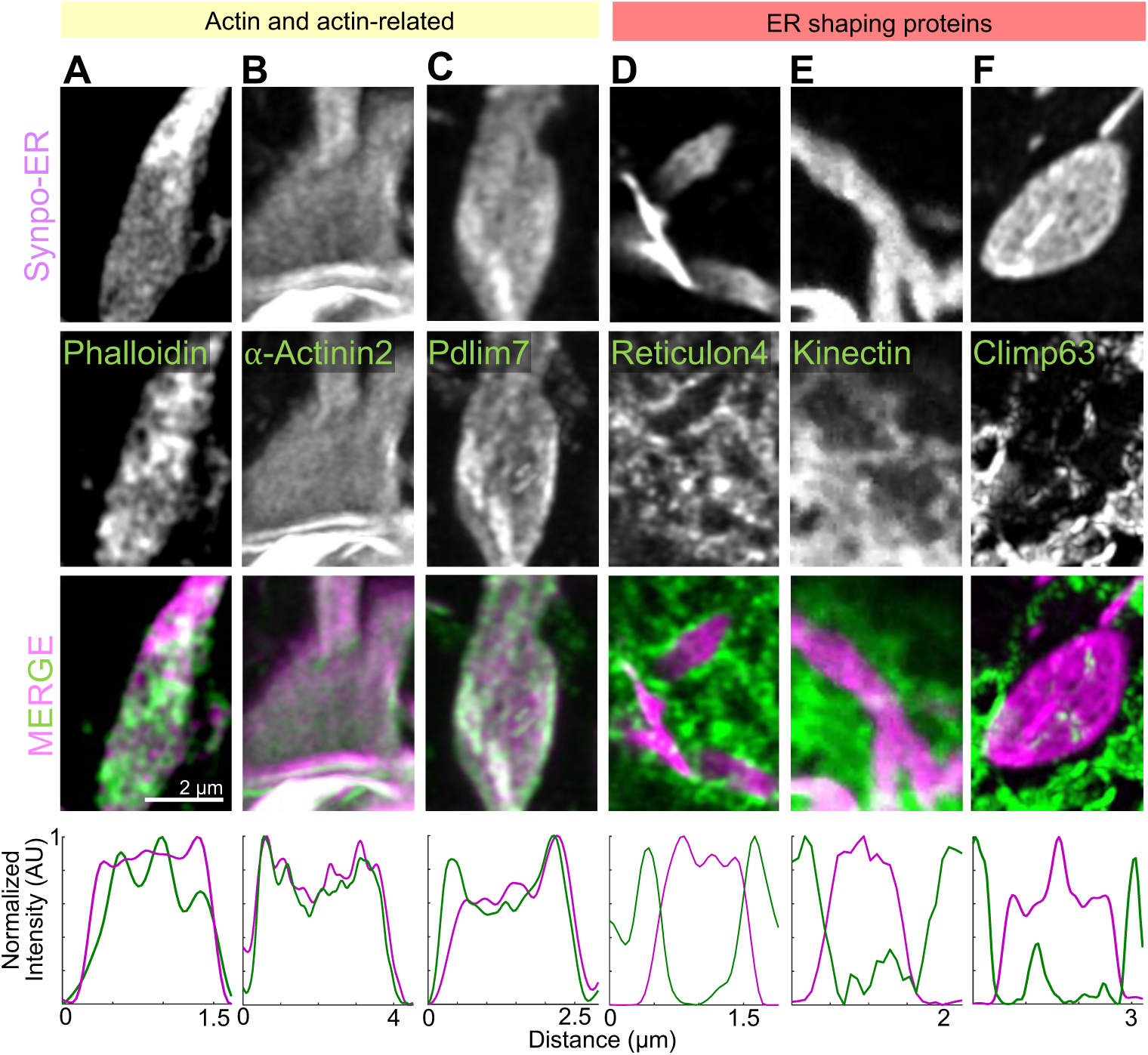
Presence of actin and actin-related proteins in SALs. **A–F**. High magnification AiryScan images of individual SALs showing the localization of Synpo-ER and of the fluorescently tagged proteins indicated. Line scan plots are shown in the bottom. Reticulon 4-GFP is enriched at the edges of SALs and excluded from their flat portions. GFP-Kinetctin and GFP-Climp63 are excluded from SALs.

### Formation of SALs is driven by a crosslinked actin meshwork on the ER surface

As SALs are represented by ER sheets, we examined the contribution to their structure of ER-shaping proteins. Reticulon 4-GFP and EGFP-Atlastin 1, which are known to localize to, and stabilize, positive curvatures in ER membranes [51–53], localized to the edges of SALs and were excluded from their flat surfaces in live COS-7 cells (Fig. 6D, S5). Lunapark, another ER shaping protein that localizes to membranes with negative curvature [54] was enriched at hot spots which corresponded to discontinuities of the synaptopodin-ER signal within the sheets (Fig. S5). These discontinuities likely correspond to the fenestrations [55, 56] exhibiting negative curvatures within the sheets as revealed by EM images of SALs (Fig. S4A blue arrow). In contrast, the two sheet-forming proteins that we investigated, namely GFP-Kinectin (Fig. 6E) and GFP-Climp63 [53, 57] (Fig. 6F), did not localize to the flat portion of SALs sheets, indicating that they do not play a role in their formation.

*En-face* views of ER cisterns of SALs stained with phalloidin revealed a dense meshwork of actin between them, supporting a model in which synaptopodin crosslinks cisterns via its multivalent binding to actin (Fig. 6A). Moreover, as discussed above, actin binding proteins that we had previously found to be components of the spine apparatus, such as Pdlim7 and *α*-actinin 2 [14], colocalized with synaptopodin-ER when co-expressed with this protein in live COS-7 cells, suggesting that they are part of the dense matrix that connect cisterns of SALs (Fig. 6B-C). In contrast, EGFP-MyosinIIA (Fig. S6A) did not accumulate at SALs, indicating that actin connecting the ER sheets is not part of an actomyosin contractile network.

The property of generating SALs if anchored permanently to the ER was not a general feature of actin binding proteins of the spine apparatus. *α*-actinin 2, an interactor of both F-actin and synaptopodin [14, 36, 42], did not show this property, as when *α*-actinin 2-GFP-Sec61*β* was expressed in COS-7 cells, its localization mirrored that of the general ER marker, DsRed-KDEL, a 27KD fluorescent protein which localizes to the ER lumen (Fig. S6C). Based on these results, we propose that the expansion and stacking of ER sheets is driven by the formation of an actin-based cytomatrix nucleated by, and bound to, synaptopodin at the interface of ER cisterns (Fig. 7).

**Fig. 7.**
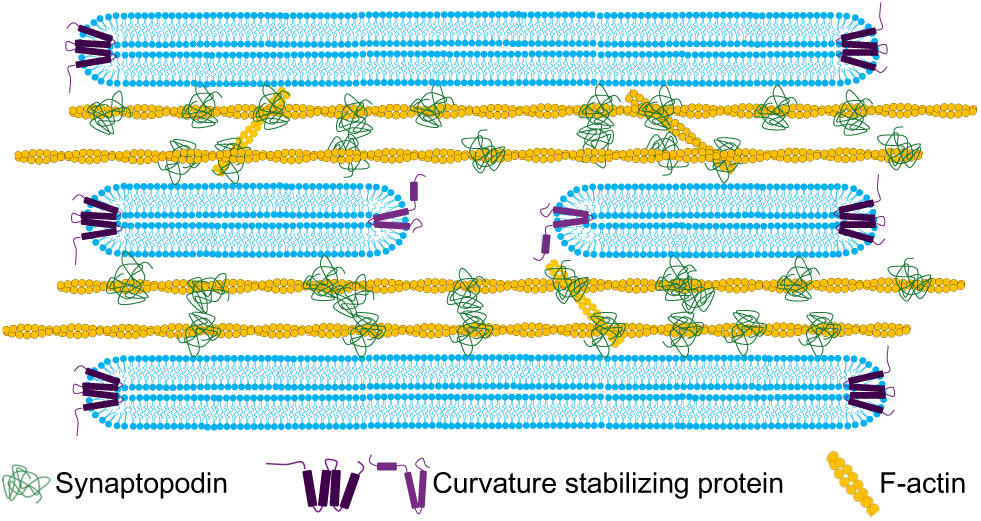
Proposed model for the formation of the spine apparatus and cisternal organelle. Synaptopodin associates with ER by an unidentified binding partner, and subsequently recruits and crosslinks an actin meshwork to this organelle. This process leads to the expansion and stacking of the ER sheets as well as to the narrowing of their lumen.

## Discussion

The spine apparatus and the cisternal organelle are two specialized subdomain of the ER that share morphological features and require the actin binding protein synaptopodin for their formation [16, 17]. In this work we show that WT synaptopodin not only can crosslink and bundle actin, as previously reported [34, 36, 42], but can also connect actin bundles to the ER in living cells, both in neurons and in non-neuronal cells. A highly conserved 45-amino-acid region within synaptopodin is necessary for its property to link actin to the ER, although this domain alone was not sufficient to target a protein to the ER.

We have further shown, using a heterologous system (COS-7 cells) that artificially anchoring synaptopodin to the ER by fusing it to Sec61*β*, which contains a C-terminal anchor [50], results in structures (SALs) that resemble the spine apparatus in key morphological features and at least some molecular properties. Like the spine apparatus, SALs consist of stacks of cisterns with a very narrow lumen connected by a dense matrix that contains actin and actin binding proteins also found in the spine apparatus of neurons. The narrow lumen is not a fixation artifact, as live imaging of SALs in cells also expressing the luminal marker, DsRed-KDEL showed absence of this marker from SALs. Known ER-sheet enriched proteins do not localize to SALs, indicating that known ER-sheet forming mechanisms do not apply to their formation and that cisterns apposition via an intervening non contractile actin-basic matrix is sufficient to expand tubules into sheets and to narrow their lumen. Contractile and non-contractile actin meshworks are known to be implicated in the dynamics of sub-cellular organelles. Our finding reveals a new role of a non-contractile actin meshwork in membrane shaping.

The distinct molecular composition of SALs, despite their continuity with the entire ER network, suggests that the spine apparatus and cisternal organelle also differ in their protein content from the rest of ER. This distinct molecular composition might create a functionally specialized subdomain within ER, tailored to meet the specific needs of the dendritic spine and axonal initial segment. The enrichment in ER membranes of these structures with minimal luminal space suggests that membrane associated functions, including lipid signaling, may predominate and that Ca^2+^ storage might not be the central function of the spine apparatus and cisternal organelle, as previously suggested. For example, IP_3_ receptor-dependent Ca^2+^ signaling is a prominent feature of the ER that populates the spine heads of Purkinje cells, but such ER elements do not form a spine apparatus.

Our results suggest that the spine apparatus and synaptopodin are conserved from flies to humans. The spine apparatus is detectable in the small subset of fly neurons that have actin-based spines resembling those found in vertebrates. This observation implies a co-evolutionary relationship between appearance of actin-based spines and occurrence of spine apparatus and is consistent with the essential role of F-actin in the formation of this organelle. Such coevolution may have provided a link between the actin-driven structural plasticity of spines and a function of the spine apparatus in regulatory mechanisms that operate within spines.

## Materials and Methods

### Antibodies and Reagents

The list of antibodies, their working dilution and their suppliers can be found in Table S1. The following constructs were kind gifts: mRFP-synaptopodin from A. Triller [Institut de Biologie de l’Ecole Normale Supérieure (IBENS), Paris, France]; EGFP-*α*-actinin-2 from J. Hell (Addgene plasmid #52669); EGFP-MyosinIIA from M. Krummel (Addgene plasmid #38297); pAcGFP-Sec61*β* from E. Schirmer (Addgene plasmid #62008); mRFP-FKBP from Tamas Balla (Addgene plasmid #67514); Reticulon 4-GFP and GFP-Climp63 from J. Bewersdorf lab (Yale). HA-EGFP-Synaptopodin, EGFP-Magi1, EGFP-Magi2, EGFP-Pdlim7, Synaptopodin-AcGFP-Sec61*β*, Synaptopodin-mCherry-Sec61*β*, pAAV-GFP-MCS, Kinectin1-EGFP, GFP-CAAX, EGFP-Atlastin1, Lunapark-mCherry, EGFP-MOSPD1, STIM1-RFP, and VAP-B-mCherry were previously constructed in our lab. AAV2/9 packaging for pAAV-HA-EGFP-synaptopodin was done by the Penn Vector Core, University of Pennsylvania.

### Generation of Plasmids

Most constructs were generated with regular PCR and/or restriction enzyme digestion and ligation. Some constructs were ligated using In-Fusion Cloning (Takara Bio). Details of primer sets, enzymes, techniques, plasmids, and cDNA clones used for each construct can be found in Table S2.

All constructs were sequenced in their entirety before use in any experiment.

### Primary Neuronal Culture and Transfection

Hippocampi of P0-P1 C57BL/6 mice (Jackson Laboratory) were dissected on ice in Hibernate-A media (Thermofisher). Dissected hippocampi were then washed in ice cold Dissociation medium (final concentration: 5.8mM MgCl_2_, 0.252mM CaCl_2_, 10mM HEPES pH = 7.4, 1mM Pyruvic Acid, 81.7% Mg and Ca^2+^ free HBSS from Thermofisher) and immediately digested in Cysteine-activated Papain solution (17U/ml Papain from Worthington, 20µg/ml DNase I from Sigma, 2mg/ml of L-Cysteine Hydrochloride from Sigma in Dissociation media) for 30 min at 37*^◦^*C. Samples were then washed in sequence with 10% FBS (in Dissociation media), Dissociation medium and Neurobasal-A medium (Thermofisher) supplemented with 2% B27 and 2mM L-Glutamax. Subsequently, 120-150K cells were plated on the glass bottom of MatTek plates coated with Poly-D-Lysine (1mg/ml) in 150µl of Neuronal Growth Medium [Neurobasal-A supplemented with 2% B27, 2mM L-Glutamax, 15% glial enriched medium (Neurobasal-A supplemented with B27 collected from a culture of DIV7+ mouse glial culture) and 10% cortical enriched medium (Neurobasal-A supplemented with B27 collected from DIV7+ mouse neuronal culture)]. 4-16hrs after plating, 2ml of Neuronal Growth Medium was added to each plate and an additional 0.5ml of Neuronal Growth Medium was added to each plate every 3-4 days afterwards. Hippocampal neurons were transfected on DIV11-13 using the CalPhos Mammalian Transfection kit (Takara) according to manufacturer’s instructions, and imaged at DIV16-28 either live or after fixation.

### Non-neuronal Cell Cultures and Transfections

COS-7 cells were obtained from ATCC and grown in DMEM (Thermo Fisher Scientific) supplemented with 10% FBS, 100 U/mL penicillin, and 100mg/mL streptomycin (Thermo Fisher Scientific) at 37*^◦^*C in humidified atmosphere at 5% CO2. They were seeded on glass bottom matTek dishes at least 16 hours prior to transfection, which was performed by Lipofectamine 2000 (Thermofisher) per manufacturers specifications. All cultured cells were routinely tested for mycoplasma contamination and found to be negative.

### Live Cell Imaging and Immunofluorescence

Confocal and AiryScan imaging were performed using LSM880, LSM800 (Carl Zeiss Microscopy) microscopes with 63X/1.40 NA plan-apochromat oil immersion objective and 32-channel gallium arsenide phosphide (GaAsP)-photomultiplier tubes (PMT) area detector. For CLEM, light microscopy was performed with an Andor DragonFly microscope with either a PlanApo 63X/1.4 NA oil immersion objective or a 20X air objective and equipped with a Zyla cMOS camera. 405nm, 488 nm, 561 nm and 633 laser lines were used.

#### Live cell imaging

Non-neuronal cells were imaged using either culturing media or Live Cell Imaging buffer (Life Technologies), and cultured neurons were imaged in modified Tyrode Buffer (119mM NaCl, 5mM KCl, 2mM CaCl_2_, 2mM MgCl_2_, 30mM glucose, 10mM HEPES, pH = 7.35).

#### Immunofluorescence

Neurons were fixed with 4% PFA, 4% sucrose, 1mM MgCl2 and 0.1mM CaCl2 in PBS. COS-7 cells were fixed in 4% PFA in PBS. In both cases fixation was carried out for 20 min at room temperature. Cells were then washed 3x with PBS and incubated sequentially in i) permeabilization buffer (0.1% saponin in PBS) for 10 mins, ii) blocking buffer (2% BSA in PBS) for 1 hour at room temperature or overnight at 4*^◦^*C and iii) with primary antibodies in blocking buffer overnight at 4*^◦^*C or phalloidin in blocking buffer for 1hr at room temperature. Samples were washed 3x in PBS and incubated with Alexa Fluor-conjugated secondary antibodies (1:500 in the blocking buffer) for an hour at room temperature followed by 3x wash in PBS.

### Image Analysis

Image analysis was performed using FIJI (RRID:SCR 002285). For measuring the distances between sheets, publicly available InteregedDistance macro by Santosh Patnaik was used. Plots were prepared using MATLAB (RRID:SCR 001622). For presentation purposes brightness and contrast of images were adjusted, and Noise *>* Despeckle function of FIJI was applied to some images. Line scan plots were generated by drawing a line across the image with a width of 10 points and using Plot Profile function of FIJI.

### Correlative Light and Electron Microscopy (CLEM)

Cells were plated on 35mm grid, glass-bottom MatTek dishes P35G-1.5-14-CGRD), infected or transfected as indicated, fixed in 2.5% glutaraldehyde in 0.1M cacodylate buffer, and imaged by light microscopy. They were subsequently postfixed in 2% OsO4 and 2% K4Fe(CN)6 (Sigma-Aldrich, St. Louis, MO) in 0.1M sodium cacodylate buffer, en bloc stained with 2% aqueous uranyl acetate, dehydrated and embedded in Embed 812. Regions of interest were sectioned (50-60nm) and imaged using a Talos L120C TEM microscope at 80kV. EM reagents are from EMS, Hatfield, PA unless noted otherwise.

## Acknowledgements

We thank members of the De Camilli lab for advice and discussions. This work was supported in part by NIH grant NS36251 and HHMI to P.D.C. We thank the Princeton FlyWire team and members of the Murthy and Seung labs at Princeton, as well as members of the Allen Institute for Brain Science, for development and maintenance of FlyWire (supported by BRAIN Initiative grants MH117815 and NS126935 to Murthy and Seung). We also acknowledge members of the Princeton FlyWire team and the FlyWire consortium for neuron proofreading and annotation.

## Declarations

- Funding: NIH grant NS36251 and HHMI to P.D.C.
- Conflict of interest/Competing interests: Authors declare no conflicts of interests.
- Ethics approval and consent to participate: Not applicable.
- Consent for publication: Not applicable.
- Data availability: All data raw data will be publicly available upon acceptance.
- Materials availability: All materials are available upon request.
- Code availability: Not applicable.
- Author contribution: P.D.C. and H.F. designed experiments. Y.W. performed EM experiments. All other experiments and analyses were performed by H.F. All authors interpreted the data. H.F. and P.D.C. wrote the manuscript with input from Y.W.

