## Supplementary Figures and Tables for "Ectopic Reconstitution of a Spine-Apparatus Like Structure Provides Insight into Mechanisms Underlying Its Formation"

### Supplementary Material

#### Supplementary Figures

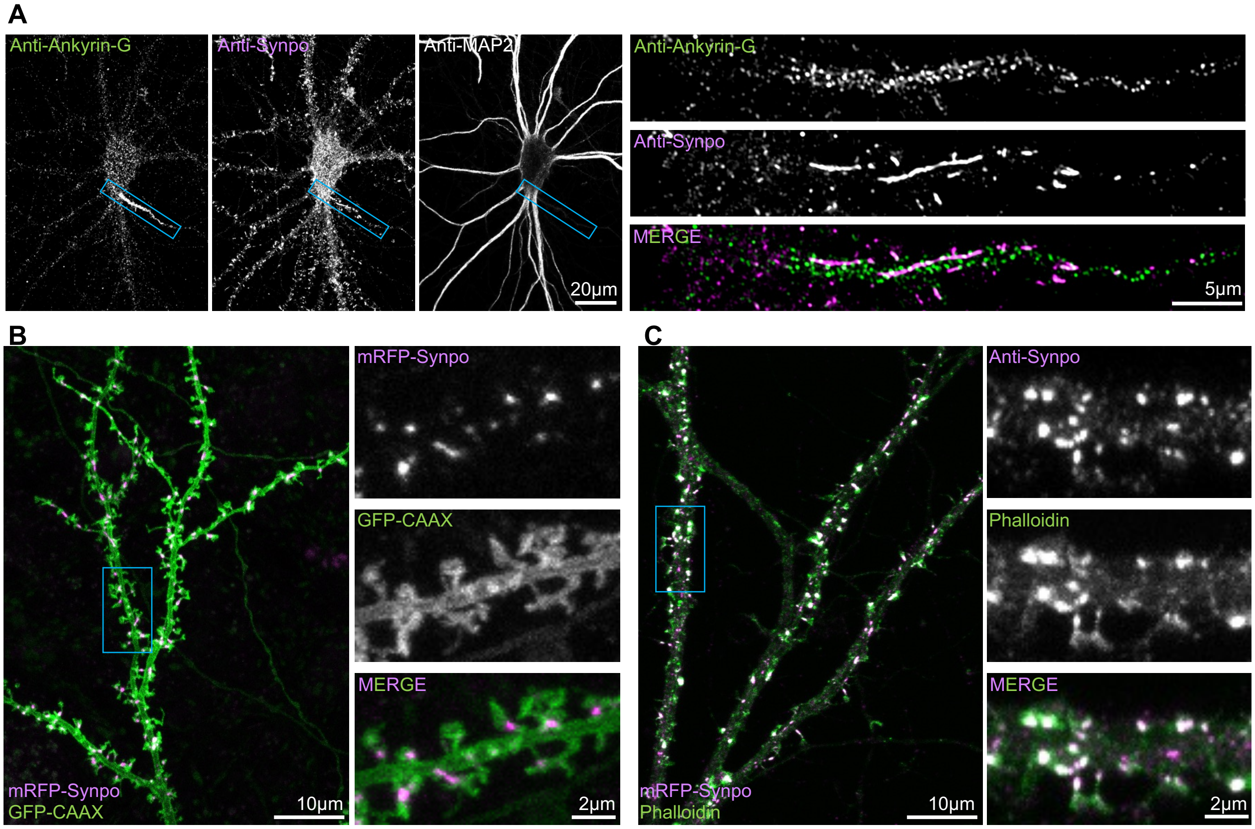


**Figure S1. Localization of synaptopodin in dendritic spines and at axonal initial segments. A.** In cultured hippocampal neurons, endogenous synaptopodin is detected by immunofluorescence in dendritic spines and at axonal initial segments. Axon initial segment (shown at higher magnification at right) is identified by its presence of immunoreactivity for Ankyrin-G (a marker of this axonal region) and absence of immunoreactivity for the dendritic marker, MAP2. **B.** Co-expression of mRFP-synaptopodin with the plasma membrane marker GFP-CAAX in a cultured hippocampal neuron reveals that synaptopodin puncta are localized at the interface between the neck and the head of dendritic spines, as demonstrated by the high magnification fields shown at right. **C.** Endogenous synaptopodin also colocalizes with a pool of F-actin in dendritic spines. Higher magnification images are shown on the right.

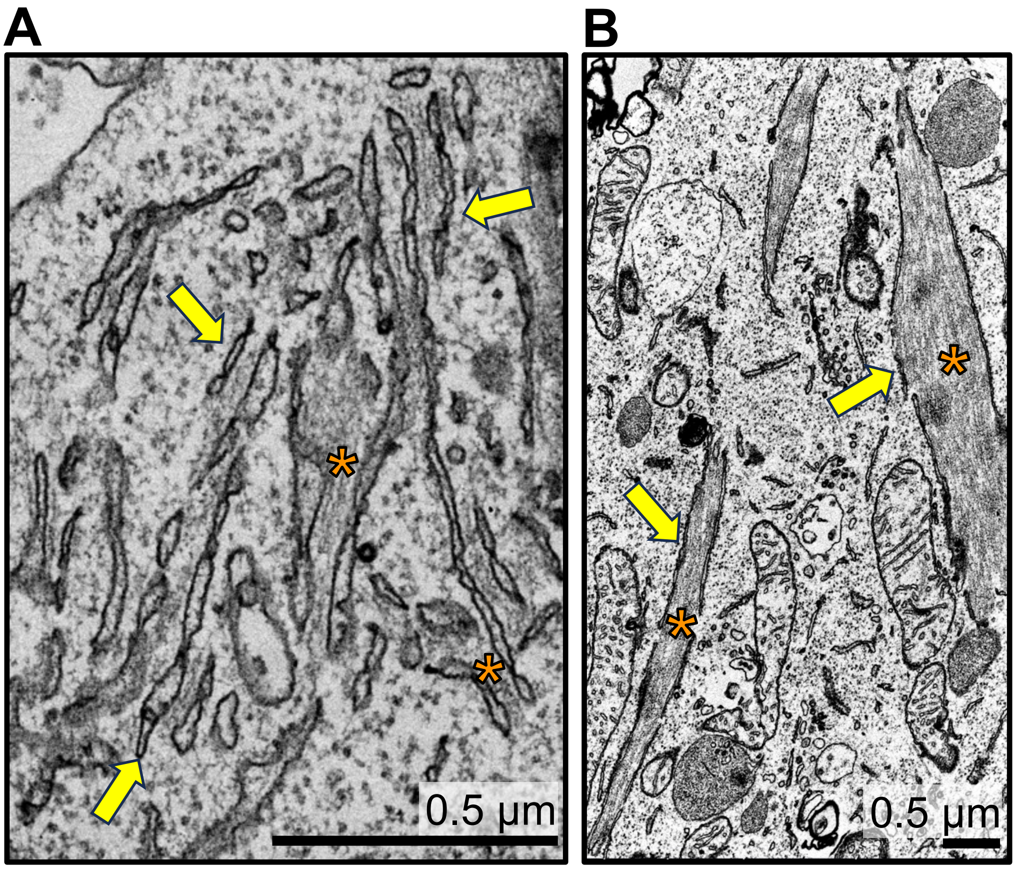


**Figure S2. Presence of actin bundles encased by ER in the soma of a cultured hippocampal neuron overexpressing synaptopodin.** Note presence of numerous actin bundles of variable thickness encased by ER.

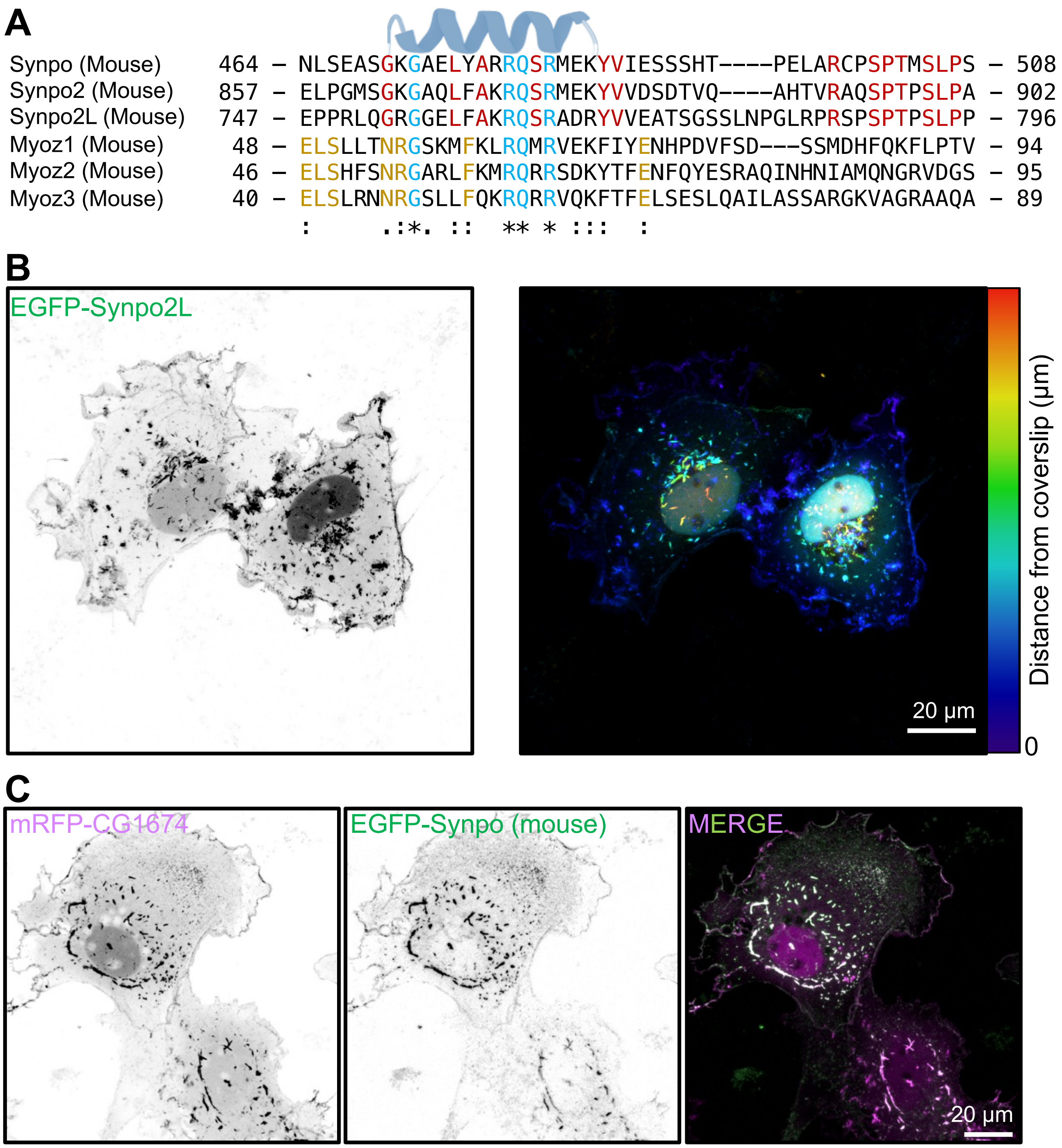


**Figure S3. Synaptopodin homologs. A.** Conservation of the calsarcin domain in synaptopodin and myozenin mouse paralogues. **B.** COS-7 cell expressing fluorescently tagged Synpo2L. The image on the right is color-coded based on the distance of the synaptopodin fluorescence from the coverslip. Similar to synaptopodin, Synpo2L localizes to plasma membrane in close proximity to the coverslip, and also forms large inclusions that can be farther away from the coverslip. **B.** COS-7 cell co-expressing the fluorescently tagged *Drosophila* orthologue of synaptopodin, mRFP-CG1674, with mouse EGFP-synaptopodin showing precise colocalization of the two proteins.

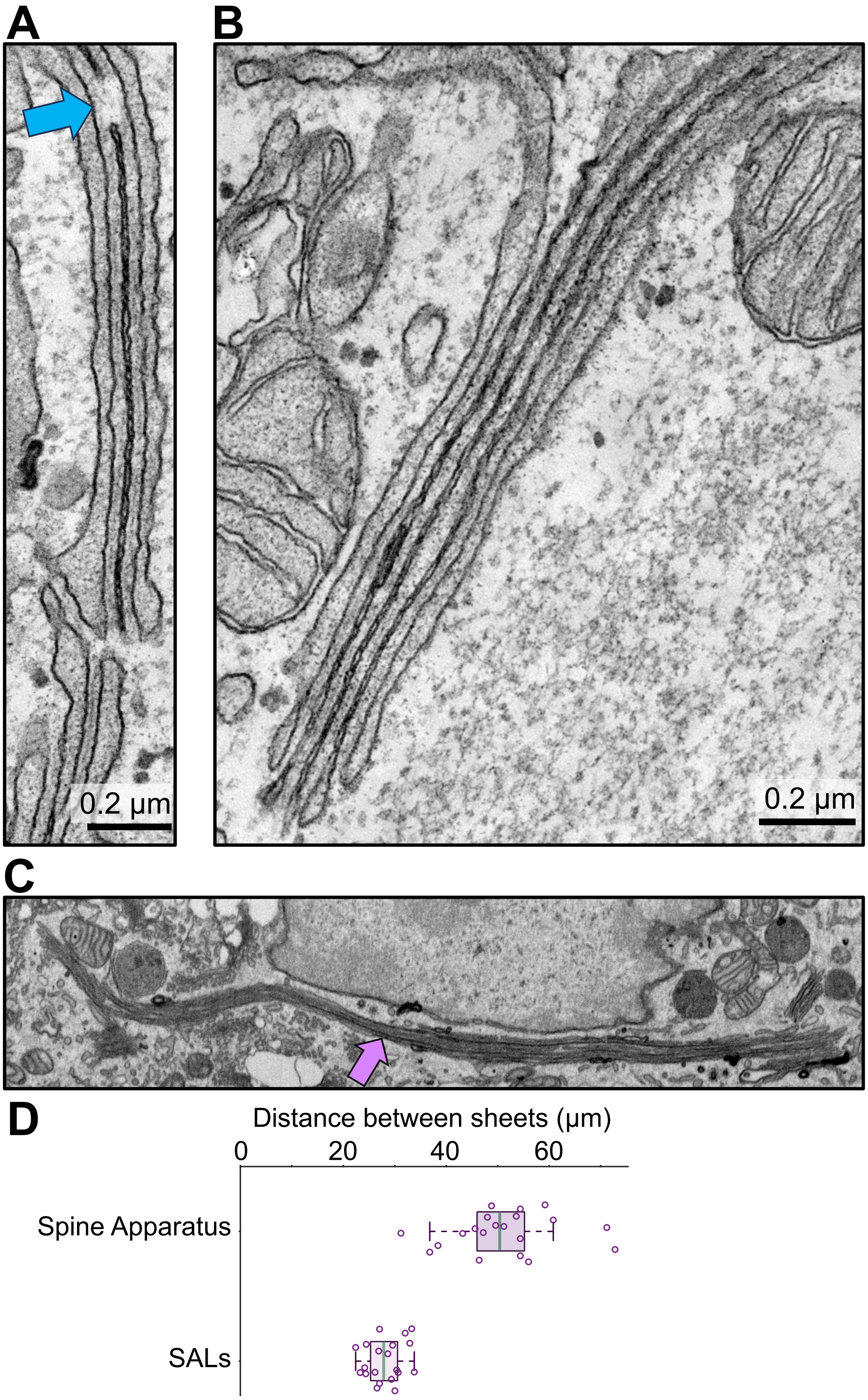


**Figure S4. Generation of SALs in COS-7 cells. A-B.** Examples of SALs induced by expressing synaptopodin-ER. A blue arrow shows a fenestration in one of the sheets. **C.** A single plane image from a stack of FIB-SEM images of a COS-7 cell expressing synaptopodin-ER. A very long SAL is indicated by a magenta arrow (see also Video 2). **D**. Quantification of the distances between ER sheets in the spine apparatus of dendritic spines and in SALs. Each circle represents the distance measurement between two opposed ER sheets in a different stack of a spine apparatus or a SAL.

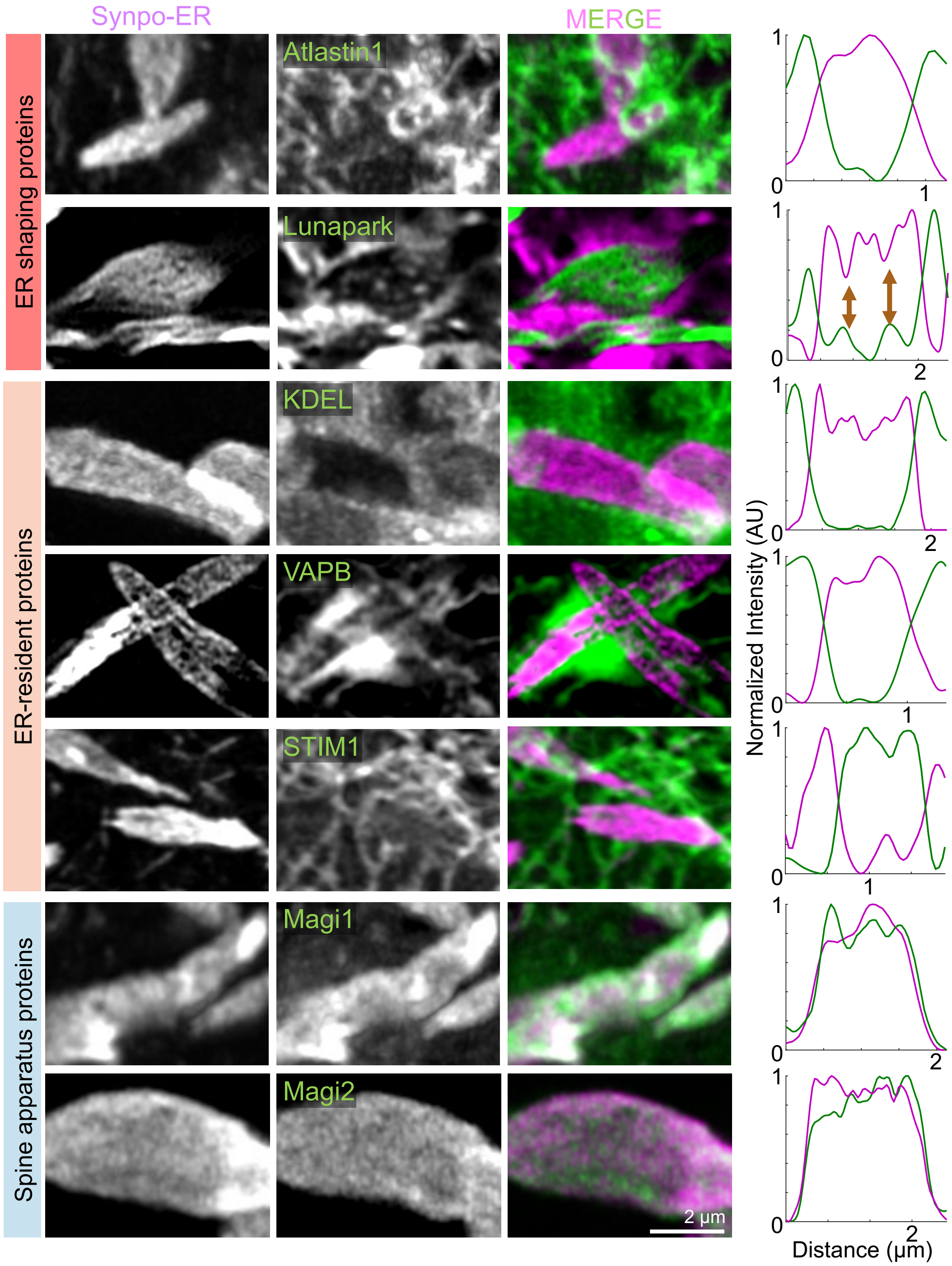


**Figure S5. Molecular characterization of SALs. A.** AiryScan images of SALs where the localization of fluorescent Synpo-ER is compared (in pairwise comparisons) to the localization of co-expressed fluorescently tagged ER shaping proteins, other ER housekeeping proteins and spine apparatus proteins. Line plots are shown on the right. The coincidence of the florescence signal of Lunapark-mCherry within the sheets of SALs, alongside discontinuities in Synpo-ER fluorescence, is indicated by gold arrows in the line plot.

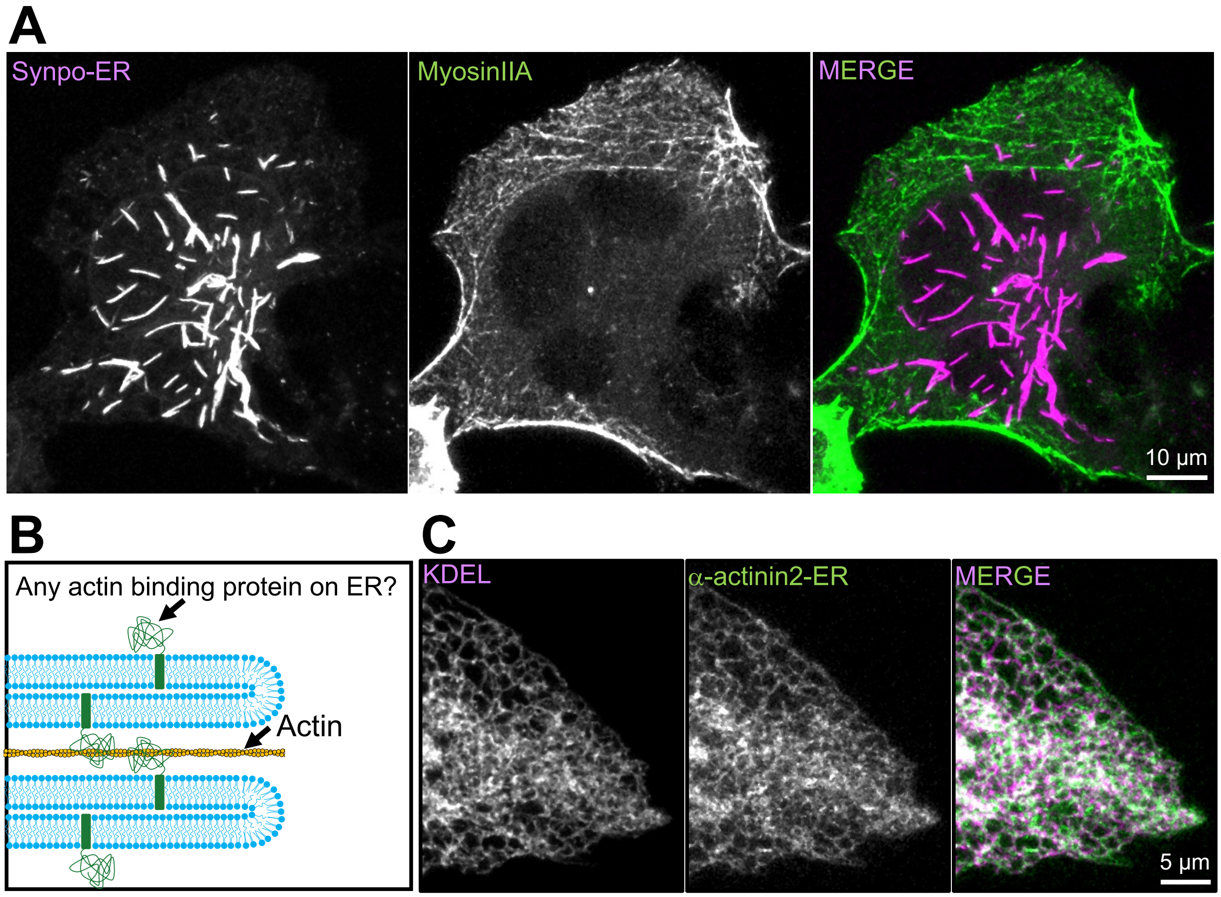


**Figure S6. Insufficiency of actin binding for induction of SAL formation. A.** COS-7 cell coexpressing Synpo-ER with MyosinIIA-GFP. **B.** Schematic of a model proposing that targeting any actin-binding protein to the ER would result in the expansion and stacking of ER sheets forming SALs. However, this model is disproven (**C**), as the targeting α-actinin2 to ER by fusing it to Sec61β does not induce SAL formation in COS-7 cells.

Supplementary tables

Table S1. List of antibodies.

| Protein (epitope) | Company; Catalog number | Antibody species | Working dilution for immunocytochemistry | Working dilution for immunoblotting |
|---|---|---|---|---|
| Synaptopodin | Sigma S9442-200UL | Rabit | 1:1000 | 1:2500 |
| Ankyrin-G | NeuroMab 75-147 Clone 106/65 | Mouse | 1:500 | N/A |
| MAP2 | Invitrogen PA1-10005 | Chicken | 1:100 | N/A |
| Phalloidin-Alexa Fluor 488 | Thermofisher A12379 | | 1:300 | NA |

Table S2. List of cloning methods and reagents.

| Construct | Cloning method | Primers | Backbone | Template/ insert |
|---|---|---|---|---|
| pAAV-HA-EGFP-Synaptopodin | Digestion / ligation with AgeI and ClaI | N/A | pAAV-BioID2-Synaptopodin | pAAV-HA-EGFP-Pdlim7 |
| mRFP-FKBP12-Synaptopodin | Digestion / In-Fusion | Primer 1: ACCGGCGCCttgtacaccggactcagatctcgaagc<br>Primer 2: agtccggacttgtacaattccagttttagaagctccacatc | mRFP-Synaptopodin | mRFP-FKBP12 |
| mRFP-Synaptopodin Δ(384-473) | In-Fusion | Primer 1: tgaccccggagctctatgcccgccgc<br>Primer 2: agagctccggggtcaccttgggcttctcc | N/A | mRFP-Synaptopodin |
| mRFP-FKBP12-Synaptopodin (1-383) | In-Fusion | Primer 1: TGACCCCGTAACACACCGCGGGCCCG<br>Primer 2: TGTGTTACGGGGTCACCTTGGGCTTCTCC | N/A | mRFP-FKBP12-Synaptopodin |
| mRFP-FKBP12-Synaptopodin (474-690) | In-Fusion | Primer 1: TCGACTTCGAGCTCTATGCCCGCCGC<br>Primer 2: AGAGCTCGAAGTCGACTGCAGAATTCGAAGC | N/A | mRFP-FKBP12-Synaptopodin |
| mRFP-FKBP12-Synaptopodin ΔCalsarcin | In-Fusion | Primer 1: CCAATCAGTCCTGGAAGTACACCACTAACGC<br>Primer 2: TCCAGGACTGATTGGGTTTGGGCTTCGG | N/A | mRFP-FKBP12-Synaptopodin |
| mRFP-FKBP12-Synaptopodin (380-690) | In-Fusion | Primer 1: tcgacttcGTGACCCCGAATCCAGATTTGC<br>Primer 2: GGGTCACGAAGTCGACTGCAGAATTCGAAGC | N/A | mRFP-FKBP12-Synaptopodin |
| mRFP-FKBP12-Synaptopodin (380-690) ΔCalsarcin | In-Fusion | Primer 1: CCAATCAGTCCTGGAAGTACACCACTAACGC<br>Primer 2: TCCAGGACTGATTGGGTTTGGGCTTCGG | N/A | mRFP-FKBP12-Synaptopodin(380-690) |
| mRFP-CG1674 | In-Fusion | For CG1674:<br>Primer 1: GCAGTCGACTTCATGGATTCTACTTTAAATATTGAGAATG<br>Primer 2: GCCCGCGGTGTGTTAAAAATCAGAGTACGGTAGATTTC<br>For Backbone:<br>Primer 1: TAACACACCGCGGGCCCG<br>Primer 2: CATGAAGTCGACTGCAGAATTCGA | mRFP-Synaptopodin | From IP15312 |
| mRFP-Synaptopodin 2L | In-Fusion | For Synaptopodin2L:<br>Primer 1: GCAGTCGACTTCATGGGTGCTGAGGAGGAGGTGC<br>Primer 2: GCCCGCGGTGTGTTACAACTGGTGCCCTGCCCC<br>For Backbone:<br>Primer 1: TAACACACCGCGGGCCCG<br>Primer 2: CATGAAGTCGACTGCAGAATTCGA | mRFP-Synaptopodin | DNASU: HsCD00861847 |
| α-actinin-2-AcGFP-Sec61β | In-Fusion | Primer 1: CGCTAGCGCTACCGGATGAACCAGATAGAGCCCGGC<br>Primer 2: CATGGTGGCGACCGGTAGATCGCTCTCCCCGTAGAG | AcGFP-Sec61β | pEGFP-α-actinin-2 |
